# Gene drive that results in addiction to a temperature sensitive version of an essential gene triggers population collapse in Drosophila

**DOI:** 10.1101/2021.07.03.451005

**Authors:** Georg Oberhofer, Tobin Ivy, Bruce A. Hay

**Affiliations:** California Institute of Technology. Division of Biology and Biological Engineering. 1200 East California Boulevard, MC156-29, Pasadena, CA 91125

**Keywords:** Gene drive, Drosophila, selfish genetic element, population suppression

## Abstract

One strategy for population suppression seeks to use gene drive to spread genes that confer conditional lethality or sterility, providing a way of combining population modification with suppression. Stimuli of potential interest could be introduced by humans, such as an otherwise benign virus or chemical, or occur naturally on a seasonal basis, such as a change in temperature. *Cleave and Rescue* (*ClvR*) selfish genetic elements use Cas9 and gRNAs to disrupt endogenous versions of an essential gene, while also including a *Rescue* version of the essential gene resistant to disruption. *ClvR* spreads by creating loss-of-function alleles of the essential gene that select against those lacking it, resulting in populations in which the *Rescue* provides the only source of essential gene function. In consequence, if function of the *Rescue*, a kind of Trojan horse now omnipresent in a population, is condition-dependent, so too will be the survival of that population. To test this idea we created a *ClvR* in *Drosophila* in which *Rescue* activity of an essential gene, *dribble*, requires splicing of a temperature-sensitive intein (TS-*ClvR^dbe^*). This element spreads to transgene fixation at 23°C, but when populations now dependent on Ts-*ClvR^dbe^* are shifted to 29°C death and sterility result in a rapid population crash. These results show that conditional population elimination can be achieved. A similar logic, in which *Rescue* activity is conditional, could also be used in HEG-based drive, and to bring about suppression and/or killing of specific individuals in response to other stimuli..

**SIGNIFICANCE STATEMENT:** Gene drive can be used to spread traits of interest through wild populations. In some contexts the goal is to suppress or eliminate the population. In principle, one way to achieve this goal is if the trait being spread confers on carriers conditional lethality in response to an environmental stimulus that is either introduced by humans into the target area at a specific time (a virus, otherwise benign chemical; a kind of species-specific insecticide), or that occurs naturally on a seasonal basis, such as a change in temperature. Here we show that *ClvR* selfish elements can be used to spread a gene that confers lethality and sterility in response to increased temperature, demonstrating that conditional population elimination can be achieved.

## Introduction

Gene drive occurs when particular genetic elements are transmitted to viable, fertile progeny at rates greater than those of competing allelic variants or other parts of the genome (reviewed in (1)). There has long been interest in the idea that selfish genetic elements mediating gene drive could be used to spread an unconditional or conditional fitness cost into a population, thereby bringing about population suppression or elimination (2–5). Selfish elements known as homing endonuclease genes (HEGs), which encode a site-specific nuclease (synthetic versions use RNA-guided nucleases such as Cas9 to achieve site-specificity), provide one approach to achieving this goal by spreading an unconditional fitness cost (6–10). Other approaches, some of which also utilize homing, seek to drive the population to an all-male state by shredding the X chromosome during spermatogenesis (11–15). Population suppression through homing can fail when homing rates are low (6, 7), and/or repair of cleaved target sites in the essential gene results in the creation of resistant alleles (c.f. (8, 9, 16)), variables that must be determined on a species- and locus-specific basis. Similar considerations apply to the use of Y-linked X shredders, which must also function when present on the highly heterochromatic Y chromosome.

An alternative approach to species-specific population suppression that does not require homing or sex ratio distortion utilizes gene drive to spread through a population (population modification) one or more transgenes that confer conditional lethality in response to a change in an environmental variable such as the presence of an otherwise benign chemical, infection with a virus, prokaryote or fungus, diapause or a change in temperature (c.f. (2, 4, 5, 17)). A central challenge with this approach is how to ensure the continued function of the (by definition) non-essential Cargo gene or genes needed to bring about conditional lethality or sterility, since loss-of-function (LOF) mutations that inactivate these components will be strongly selected for. An approach that eliminates the possibility of transgene inactivating mutations resulting in loss of condition-dependent lethality, and that we implement here, uses gene drive to make the survival of individuals under permissive conditions – as a necessary consequence of gene drive-based population modification – dependent on the activity of an essential gene engineered to lack function under non-permissive conditions.

### *Cleave and Rescue* (*ClvR*) selfish genetic elements as a tool for temperature sensitive population suppression

To achieve these ends, we sought to develop condition-dependent versions of the *Cleave and Rescue* (*ClvR*) selfish genetic element (18, 19) (also referred to as toxin antidote recessive embryo (TARE) in a related proof-of-principle implementation (20)). *ClvR* has two components. The first is a DNA sequence modifying enzyme such as Cas9 and one or more gRNAs. These constitute the *Cleaver*, are expressed in the germline and act in *trans* to disrupt the endogenous version of an essential gene, creating potentially lethal LOF alleles in the germline, and in the zygote due to maternal carryover of active Cas9/gRNA complexes. The second is a recoded version of the essential gene resistant to cleavage that acts in *cis* to guarantee the survival of those who carry it (the *Rescue*). The lethal LOF phenotype manifests itself in those who fail to inherit *ClvR* and have no other functional copies of the essential gene, while those who inherit *ClvR* and its associated *Rescue* survive. In this way, as with many other toxin-antidote-based selfish genetic elements found in nature (reviewed in (21)) and created de novo (22), *ClvR* gains a relative transmission advantage that can drive it to transgene or allele fixation by causing the death of those who lack it (18–20, 23). Importantly, once a *ClvR* element has spread to transgene fixation (and unlike other selfish elements in Nature), all endogenous wild-type alleles of the essential gene have been eliminated through cleavage and LOF allele creation. At this point the only source of essential gene function comes from *ClvR* itself—a form of genetic addiction—creating a state of permanent transgene fixation. In consequence, if function of the *Rescue*, a kind of Trojan horse now omnipresent in a population, is condition-dependent, so too will be the survival of that population.

One environmental cue that could in principle be used to bring about conditional lethality associated with a population crash is seasonal temperature. *Drosophila suzukii*, an invasive species of Europe, Asia and North and South America (24, 25), is one potential target for such an approach. It has a number of generations per year and is often invasive in temperate climates that experience large seasonal temperature variations (26), providing opportunities for introducing a temperature-dependent population bottleneck as a method of suppression. As a proof-of-principle demonstration of this idea we sought to create a version of *ClvR* in *Drosophila melanogaster* in which *Rescue* function is temperature sensitive (TS; TS-*ClvR*). We show that a TS-*ClvR* element can successfully spread a conditional *Rescue* into *Drosophila* populations. When populations now dependent on this transgene are shifted to non-permissive temperatures, they rapidly become sterile and go extinct.

## Results

### Insertion of a TS-intein into the *Drosophila* essential gene *dribble* (*dbe*) results in temperature-sensitive loss of function

Traditional approaches to generation of dominant or recessive TS mutations in essential genes in metazoans are laborious as they involve random mutagenesis of whole genomes followed by large-scale screens at different temperatures for otherwise fit TS mutants. As an alternative we sought to create TS versions of an essential gene by introducing a TS version of an intein into the protein coding sequences of *Rescue* transgenes within *ClvR*s previously shown to spread into wildtype populations (Fig. 1 and (18, 19)). An intein is a protein-encoded autoprocessing domain able to excise itself from a polypeptide and rejoin the N-and C-terminal flanking sequences (exteins) to create a WT version of the encoded protein (27). Importantly, once an intein has been introduced into the coding sequence of an essential gene and that version provides the only source of essential gene function, splicing activity cannot be lost through mutation since the non-spliced version is non-functional.

**Fig. 1.**
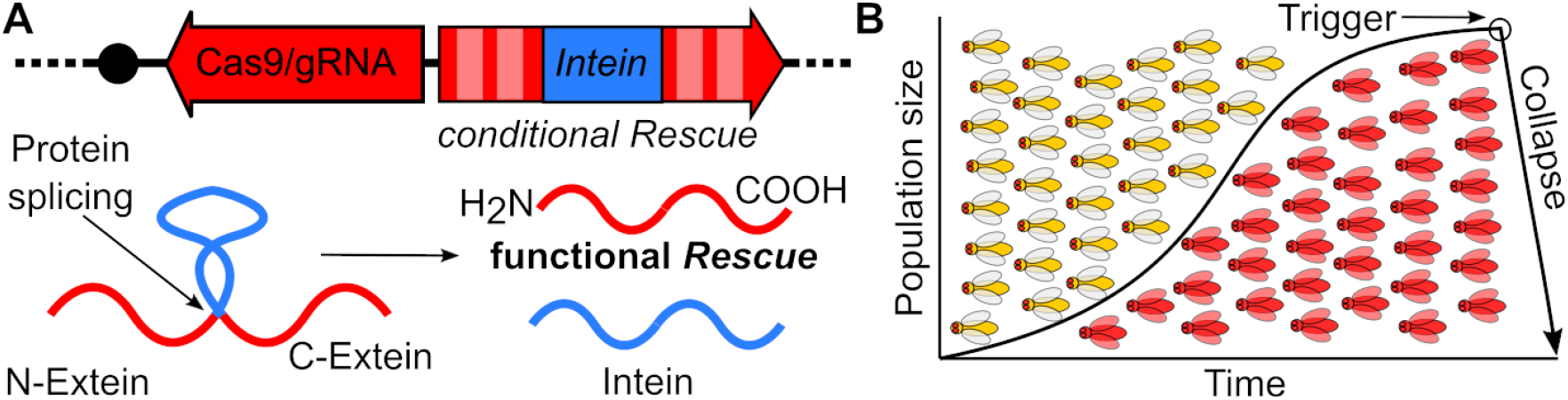
TS-*ClvR* design and concept. **(A)** TS-*ClvR* drive element comprised of Cas9/gRNAs targeting an essential gene and a recoded *Rescue* of that gene with a TS-intein within its coding region. After translation the TS-intein can splice itself out to yield a functional *Rescue* protein. **(B) Population suppression with a TS-*ClvR*.** TS-*ClvR* bearing flies (red) are released into a WT population (yellow). The TS-*ClvR* selfish element spreads into the population at the cost of WT. Once the TS-*ClvR* element has reached genotype fixation (has at least one Copy of TS-*ClvR*) in the population, all functional endogenous copy of the essential gene targeted by TS-*ClvR* will have been mutated to LOF. At this point the *conditional* TS-*Rescue* within the *ClvR* element provides the only source of essential gene function in the population, making it subject to a collapse in response to a temperature shift.

The *Sce* VMA intein, which is located within the *Saccharomyces cerevisiae* vacuolar membrane ATPase, is able to excise itself from a number of foreign proteins (28). TS versions of *Sce* VMA inteins have been isolated that allow spicing at a range of low, but not higher temperatures (ranging from 18°C to 30°C (29, 30)). A mechanistic requirement for successful intein splicing is that the C-terminal extein starts with a cysteine residue. Other less well characterized sequence contexts also regulate splicing efficiency (31–33). To determine if *ClvR Rescue* genes that contain the *Sce* VMA intein are functional we generated twelve WT- and TS-intein-bearing versions of *Rescue* transgenes for two previously described *ClvR* target genes, (*dribble* [*dbe*, in *ClvR^dbe^* (19) and *technical knockout* [*tko*], in *ClvR^tko^* (18), Fig. S1). We tested the ability of intein-bearing *Rescue* transgenes to provide essential gene function by examining progeny of a cross between females heterozygous for complete *ClvR^dbe^* or *ClvR^tko^* elements and males heterozygous for the corresponding WT-intein *Rescue* (*Rescue*-INT^WT^) or TS-intein *Rescue* (*Rescue-*INT^TS^) transgene.

When present in females, *ClvR^dbe^* and *ClvR^tko^* cleave and create LOF alleles of their target genes in the maternal germline and the zygote with a frequency of >99.9%. Thus, in the absence of another source of *Rescue* activity essentially all viable progeny should be *ClvR*-bearing (in an outcross the 50% that fail to inherit *ClvR* die because they lack a functional copy of the essential gene). In contrast, if the *Rescue*-INT^WT^ or *Rescue*-INT^TS^ in heterozygous males is active, ~33% of viable progeny should be non-*ClvR*-bearing, and these should all carry the intein-bearing *Rescue*. From crosses carried out at 23° C and 27° C we identified one version of the *dbe Rescue* that retained function, in which the intein was inserted N-terminal to cysteine 2 of the *dbe* coding sequence (Table S1 and S2). The *dbe Rescue* transgene carrying the WT-intein was functional at 23° C and 27° C. The *Rescue* carrying the TS-intein was also functional at 23°C but was largely (though not completely) non-functional at 27°C (see Fig. 2 and Table S2). Flies carrying the *dbe Rescue-*INT^TS^ construct were then used as a genetic background in which to create flies carrying a full *ClvR^dbe^*-INT^TS^ (referred to as TS-*ClvR^dbe^*) drive element carrying the other components found in *ClvR^dbe^* (19). These include Cas9 expressed under the control of the germline regulatory sequences from the *nanos* gene, four gRNAs targeting the endogenous *dbe* locus expressed under the control of individual U6 promoters, and an *OpIE-td-tomato* marker gene (Fig. S1B,C).

**Fig. 2:**
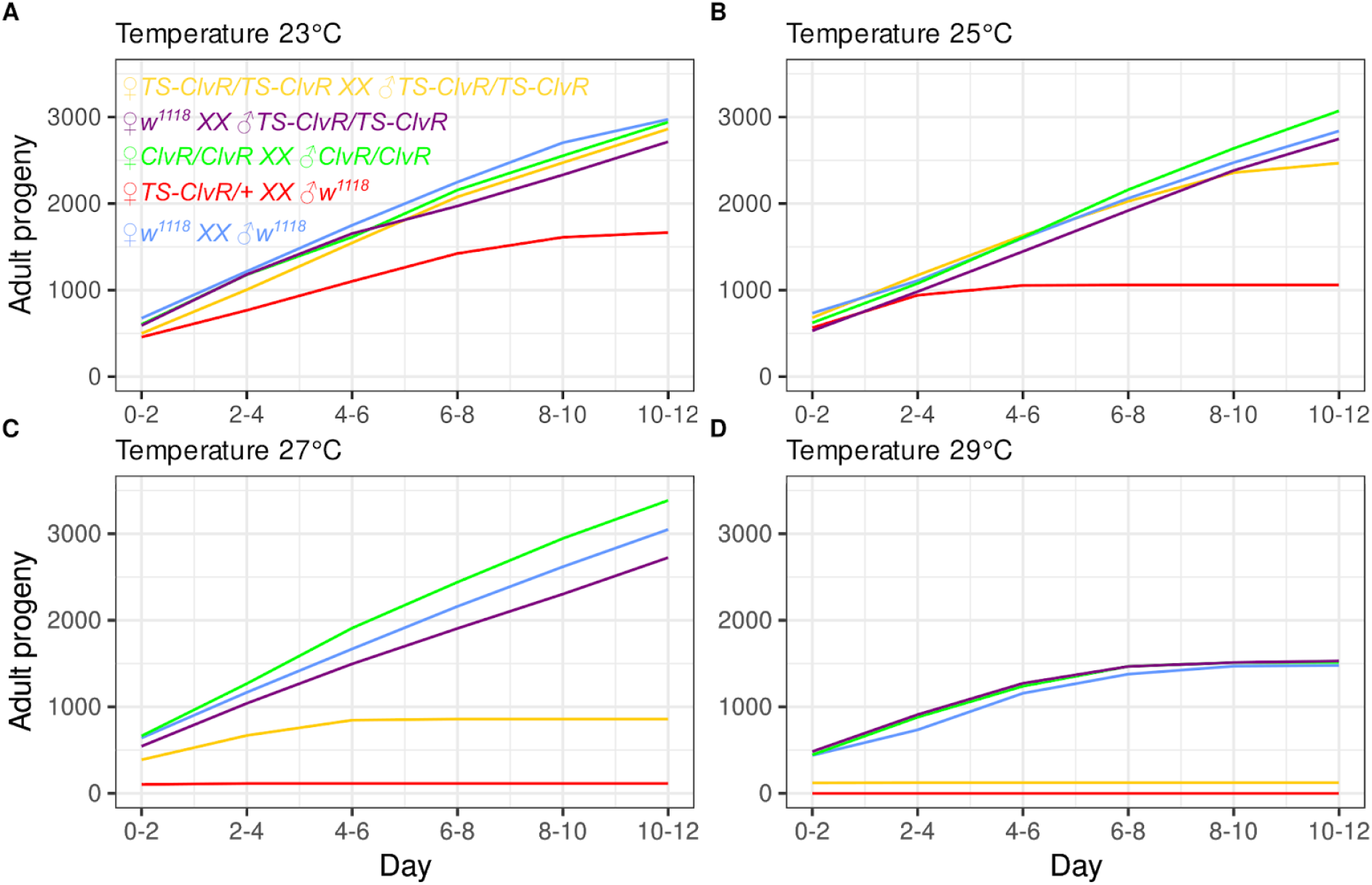
Cumulative adult fly output at different temperatures. Shown is the cumulative adult progeny output of four replicates in which 5 females were crossed to 5 males over 12 days. Crosses were heterozygous ♀*TS-ClvR^dbe^/+ XX ♂w^1118^* in red, homozygous ♀*TS-ClvR^dbe^/TS-ClvR^dbe^ XX ♂TS-ClvR^dbe^/TS-ClvR^dbe^* in yellow, ♀*w^1118^ XX ♂TS-ClvR^dbe^/TS-ClvR^dbe^* in violet, ♀*w^1118^XX ♂w^1118^* (control) in blue, and the original non-*TS* ♀*ClvR^dbe^ XX ♂ClvR^dbe^*(control) in green.

### TS-*ClvR^dbe^* efficiently creates LOF alleles at permissive temperatures

A TS-*ClvR* must be able to efficiently create LOF alleles at all relevant environmental temperatures, and Cas9 activity has been shown to be temperature sensitive, with reduced activity at lower temperatures (34, 35). To test the ability of Cas9 to create *dbe* LOF alleles at temperatures permissive for intein splicing we crossed heterozygous TS-*ClvR^dbe^* females to *w^1118^* (WT) males at 22°C and scored viable progeny for inheritance of the TS-*ClvR^dbe^* marker. As discussed above, if the TS-*ClvR^dbe^* Cas9/gRNAs successfully create *dbe* LOF alleles in the maternal germline and in the early embryo, viable progeny should be largely or exclusively TS-*ClvR^dbe^-bearing. ClvR* was present in 93.8% of the offspring, a lower frequency than previously reported for the original *ClvR^dbe^* (>99% (19)), in which crosses were carried out at 26°C. This is likely due to reduced Cas9 activity since similar tests with the original *ClvR^dbe^* stock at 22° C also resulted in a reduced drive inheritance of 95.9% (Table S3). In any case, the results of crosses, and sequencing of genomic DNA of escapers from the above crosses, show that the modestly reduced rate of cleavage was not associated with the creation of functional, cleavage resistant alleles (Data S1).

### Female TS-*ClvR^dbe^* flies suffer a temperature-dependent loss of reproductive output

In order to bring about condition-dependent population suppression following gene drive-based population modification, carriers must experience a high fitness cost under non-permissive conditions. A major determinant of fitness is reproductive output, which requires ongoing adult germline and somatic cell proliferation and growth. *Dbe* is a gene whose product is required in all proliferative cells (36). Thus, reproductive output is likely to be a sensitive indicator of *dbe* function and the effects of dosage at different temperatures. To explore these topics, we characterized the reproductive output of females having two, one or no copies of TS-*ClvR^dbe^*. We focused on females because adult sexual maturation requires cell proliferation and growth of somatic and germline cells. In contrast, young adult males already contain large numbers of mature sperm, which have a long functional lifetime once deposited in the female reproductive tract (37). For each cross, four replicate vials having 5 females and 5 males (derived from flies raised at 22°C) were incubated at different temperatures ranging from 23° C to 29° C, and transferred to fresh vials every two days. The cumulative adult fly output from these crosses over time is plotted in Fig. 2 (see also Fig. S2). At the low temperature of 23° C, crosses between homozygous WT (*w^1118^*) flies resulted in the production of progeny at a roughly constant rate, with only a modest drop off in production during days 10-12. The rate of offspring production over time was similar for crosses involving homozygous (non-TS) *ClvR^dbe^* males and females, and for crosses between WT females and homozygous TS-*ClvR^dbe^* males (both *ClvR*s were created in a *w^1118^* genetic background). In contrast, crosses between heterozygous TS-*ClvR^dbe^* females and WT males produced fewer absolute numbers of progeny. This is expected since the ~50% of progeny that fail to inherit TS-*ClvR^dbe^* die due to lack of essential gene function. More importantly, the rate of offspring production also decreased significantly over time, suggesting that in an otherwise LOF background, even at permissive temperatures, one maternal copy of the *dbe Rescue*INT^TS^ results in gradual loss of *dbe*-dependent maternal germline activity required for reproduction.

At higher temperatures (25°C-27°C) the loss of reproductive potential of TS-*ClvR^dbe^*-bearing adult females as compared to WT or those carrying *ClvR^dbe^* was more dramatic. At 29°C heterozygous TS-*ClvR^dbe^* females became sterile immediately, while homozygous TS-*ClvR^dbe^* flies became sterile after 2 days. Progeny production also ended somewhat prematurely at 29°C for crosses in which the female parent was WT or *ClvR^dbe^-bearing.* However, this appears to be a general temperature effect since the ability to produce progeny was lost at a similar rate for both sets of crosses. These results, along with those described above involving crosses of *ClvR^dbe^* /+ females to *dbe Rescue*INT^TS^ males at different temperatures, and data presented in Tables S3 and S4, show that females carrying TS-*ClvR^dbe^* (the vast majority of which lack *dbe* function from the endogenous locus in the germline and early embryo; Table S3) are reproductively fit at lower temperature, but rapidly lose the ability to reproduce at elevated temperatures.

### TS-*ClvR^dbe^* spreads to transgene fixation at a permissive temperature

Population modification followed by suppression requires that drive into a WT population succeed at low, permissive temperatures. To test the ability of TS-*ClvR^dbe^* to achieve this end we carried out a gene drive experiment at 22° C. To seed the drive, we crossed heterozygous TS-*ClvR^dbe^* males (*w^1118^*; TS-*ClvR^dbe^*/+) to WT (*w^1118^*) females to create a starting TS-*ClvR^dbe^* allele frequency of 25%, in four replicate populations. Mated females were allowed to lay eggs in a food bottle for one day and removed afterwards. The drive experiments were kept in a temperature-controlled incubator at 22° C. After ~16 days most progeny had developed into adults, which were then removed from the bottles, scored for the presence of the TS-*ClvR^dbe^* marker (*td-tomato*), and transferred to a fresh food bottle to repeat the cycle. Results of the drive experiment are shown in Fig. 3A. The TS-*ClvR^dbe^* construct reached genotype fixation between 9 and 10 generations in all 4 replicate drive populations, while a construct carrying only the *dbe Rescue-*INT^TS^ but no Cas9/gRNAs did not increase in frequency. By generation 18 TS-*ClvR^dbe^* allele frequencies ranged from 93.2-97.6% (Table S5).

**Fig. 3.**
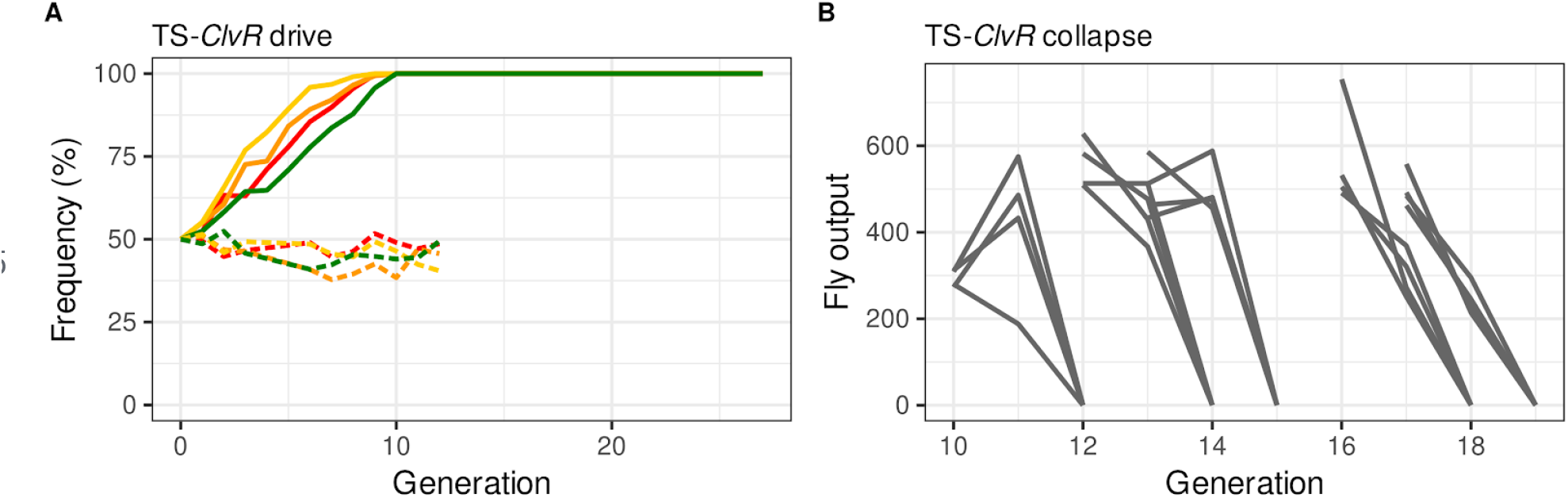
Population modification at a permissive temperature followed by suppression at a restrictive temperature. **(A)** Shown are genotype frequencies of TS-*ClvR^dbe^*-bearing flies over discrete generations at 22°C. TS-*ClvR^dbe^* is indicated with solid lines, *dbe Rescue-*INT^TS^ controls with dashed lines. **(B)** Gray lines show individual population trajectories for all replicates when incubated at 29°C. All populations produced some offspring when moved from 22°C to 29°C. These collapsed in the next generation due to complete sterility.

### Populations in which TS-*ClvR^dbe^* is ubiquitous undergo a population collapse when shifted to elevated temperature

The goal of drive with a TS-*ClvR* is ultimately to bring about a population crash in response to an environmental temperature shift once LOF allele creation associated with population modification has rendered all members of the population dependent on the *Rescue*-INT^TS^. As a test of this hypothesis, we followed the fate of drive populations shifted to 29°C at generations 10, 12, 13, 16 and 17. At each of these points adults from the 22°C drive population were allowed to lay eggs for one day at 22°C in order to continue the drive, and then moved to 29°C to allow egg laying for a further two days. Adults were then removed and the fate of the 29°C populations followed, as with the drive populations kept at 22°C (Table S6). Populations fixed for *ClvR^dbe^* (control) individuals produce many adult progeny over 6 generations when continuously housed at 29°C (c.f. Table S7). In contrast, populations of drive individuals–which at this point are heterozygous or homozygous for TS-*ClvR^dbe^*–give rise to only a few adult progeny per parent for one more generation (c.f. gray line leading from the number of generation 10 individuals transferred to 29°C to the generation 11 adult progeny number). These latter adults were universally sterile, resulting in population extinction in the next generation (Fig. 3D).

## DISCUSSION

Our results show that gene drive can be used to spread a trait conferring conditional lethality into an insect population, resulting in a population crash when the restrictive condition, in this case a temperature shift, is experienced. Additional Cargo genes, designed to bring about some other phenotype such as disease suppression prior to temperature-dependent population suppression could also be included in such gene drive elements. The implementation described herein used the *ClvR* gene drive mechanism, which concurrently renders LOF endogenous copies of an essential gene and replaces them with a TS version as spread occurs. A similar outcome (drive followed by condition-dependent suppression) could also be achieved using strategies in which a HEG homes into an essential gene locus, thereby disrupting its function, while also carrying a cleavage-resistant version of the essential gene as a rescuing transgene (38–42), that in this case is engineered to be temperature sensitive.

Conditional populations suppression systems target both males and females when a sex-independent essential gene is utilized for cleavage and conditional rescue, as described here. With such a system the target environment may require some level of periodic repopulation with transgenes. A modified system that would reduce this need, and work to maintain the transgene in the target environment in the face of incoming migration of WT, eliminates only females or female fertility under non-permissive conditions (for modeling of a related system with these characteristics see (43)). *ClvRs* that bring about LOF and *Rescue* of two different genes, one that is needed for sex-independent viability (mediating strong drive) and a second that is required for female viability or fertility (allowing for elimination of females under non-permissive conditions), could be used to achieve this goal. *ClvR*s able to rescue the viability and fertility associated with LOF of two different essential genes at the same time have been created (18, 19)), suggesting this approach is plausible. Finally, we note that the strategy for generating TS strains described here (replacement of a WT version of an endogenous gene with a TS-version) could also be used as a method of sex-specific sorting in inundative suppression strategies such as the sterile insect technique.

Success with any TS gene drive system in the wild will require knowledge of temperature fluctuations within a season in the region of interest, the life phases in which the target species is most susceptible (and resistant) to loss of essential gene function, and potentially further selections in rapidly reproducing organisms like yeast (29, 30) for TS-inteins best suited to the environmental temperature regimes involved. Also, because seasonal temperatures do not change in an all or none fashion, gradual shifts towards non-permissive conditions will provide opportunities for selection to take place on sequences within the intein coding region that reduce or eliminate temperature sensitivity. The targeting of biosynthetic essential genes such as *dbe*, whose transient LOF is unlikely to result in an immediate fitness cost (as is seen for some other TS mutants that cause immediate paralysis; c.f. (44)) probably provides some level of environmental phenotypic buffering in this regard but would not eliminate selection. While next generation *ClvR* elements can be cycled through a population, replacing old, failed elements with new ones (19), strategies that forestall the need for such cycles of modification for as long as possible would be useful. This can be achieved by building into the *Rescue* transgene mechanistic redundancy with respect to how temperature sensitivity is achieved, thereby necessitating multiple mutational hits for the *Rescue* to lose its TS characteristic. As an example, an N-terminal TS degron (the N-terminal location preventing the loss of degron activity through frameshift or stop codons) that promotes the degradation of a linked C-terminal protein at elevated temperature provides one such approach (45). Insertion of multiple copies of a common TS intein at different positions provides another.

Finally, we note that a similar logic to that presented here, in which *Rescue* activity is conditionally blocked, could be used to bring about species-specific suppression in response to other stimuli. Small molecules provide one example. These could block intein splicing activity (46), promote the degradation of a target protein (47), or decrease the stability of specific transcripts (48). Target genes that might be particularly amenable to such approaches, which will likely alter expression only transiently following application, include those encoding proteins whose loss results in rapid cell death, such as inhibitors of apoptosis (49). Virus infection provides a further opportunity for engineering conditional lethality. As an example, virus-encoded protease activity, required for viral polyprotein processing in many systems, serves as an “honest” and specific indicator of infection. If one or more viral protease target sites are engineered into the products of key host essential genes–-and these versions replace WT counterparts during drive–-cleavage at these sites in organisms that are virally infected could result in a lethal LOF phenotype. This could be used to directly suppress populations in response to introduction of a naturally-occurring and otherwise benign virus. A similar strategy could also be used to selectively eliminate members of a disease vector population that are infected with a human, animal or plant pathogenic virus, in the context of a simple population modification scenario.

## Acknowledgments

Stocks obtained from the Bloomington Drosophila Stock Center (NIH P40OD018537) were used in this study.

## Author Contributions

Conceptualization, G.O., T.I. and B.A.H.; Methodology, G.O., T.I. and B.A.H.; Investigation, G.O. and B.A.H.; Writing – Original Draft, G.O. and B.A.H.; Writing – Review & Editing, G.O., T.I. and B.A.H.; Funding Acquisition, G.O. and B.A.H.

## Data availability

All data is available in the main text or the supplementary materials.

## Supplementary Materials for

### Materials and Methods

#### Synthesis of TS-*Rescues* for *tko* and *dbe* target genes

All constructs in this work were assembled with Gibson cloning (50). Enzymes were from NEB, cloning and DNA extraction kits from Zymo. Inteins were gene synthesized as gblocks from IDT. We started from our previously cloned *Rescue* constructs (18, 19). The *Rescue* for *tko* was derived from the ortholog of *Drosophila virilis*, the one for *dbe* from *Drosophila suzukii*, Both genes have 3 cysteines in their coding sequences. We used Gibson assembly to insert a WT-intein and a TS-intein (mutation D324G; (29, 30)) after each of the cysteines for a total of 12 constructs. In addition, the plasmids had a dominant *OpIE*-GFP marker, an attP site, and homology arms to facilitate CRISPR-HR mediated insertion into the fly genome at the 68E map position on chromosome 3.

The constructs were injected into *w^1118^* flies along with a pre-loaded Cas9/gRNA RNP complex having a gRNA (both from IDT) targeting chromosome 3 at 68E (Fig. S1A). Details were as described previously (19). All Gibson cloning primers and construct Genbank files are in Data S1. Embryonic injections were carried out by Rainbow Transgenic Flies (Camarillo, USA). Injected G0 flies were outcrossed to *w^1118^* and screened for ubiquitous GFP expression.

#### Screening crosses for temperature-dependent *Rescue* activity

To determine if any of the intein-bearing *Rescues* showed temperature-dependent *Rescue* activity we set up crosses between heterozygous virgins that carry the original non-TS *ClvR* element and heterozygous males carrying the different *Rescue*-INT^(TS or WT)^ versions (Crossing scheme in Fig. S3). All crosses were set up in triplicates and incubated at 23° C or at 27° C. None of the intein-*Rescues* for *tko* were able to provide adequate gene function at either temperature (Table S1). For *dbe* the *Rescue* transgenes carrying the *WT-intein* inserted after cysteine 2 and 3 were able to rescue flies at both temperatures. *Rescue* transgenes containing the TS-intein inserted after cysteines 1 or 3 were not able to provide *Rescue* function at either temperature. In contrast, *Rescue* transgenes carrying the TS-intein inserted after cysteine 2 showed promising behavior, with most progeny dying at 27° C but not at 23° C (Table S2, highlighted in red). We used these flies to build a fully functional TS-*ClvR* selfish element. Note: For the WT-intein inserted after cysteine 1 of *dbe* we did not obtain transformants after a first round of injections. Since the TS-intein version of that construct did not show *Rescue* activity, this insertion position was not further pursued.

#### Synthesis of TS-*ClvR^dbe^* flies

Cas9 and a set of 4 gRNAs (each driven by a U6 promoter) that target endogenous alleles of *dbe* were integrated into the attP site within the TS-intein *Rescue* construct, as described previously (18, 19). The gRNA scaffolds were optimized as described previously by replacing the T base at position 4 with a G and extending the duplex by 5 bp (51, 52).

The construct was modified further using Gibson assembly to add in a new *OpIE-td-tomato* marker gene (the original plasmid had a *3xP3*-GFP marker that would have been hard to screen for in the ubiquitous GFP background of the TS-*Rescue* carrying flies) and was injected into flies carrying the TS-*Rescue* alongside a helper plasmid providing a source of PhiC31 integrase (Rainbow Transgenic Flies) (Fig. S1B). Injected G0 flies were outcrossed to *w^1118^* and screened for ubiquitous *td-tomato* expression. Positive transformants were balanced over TM3, *Sb* to subsequently generate a homozygous stock of TS-*ClvR^dbe^* flies carrying the TS-*Rescue* and Cas9/gRNAs (Fig. S1C). Primers and construct Genbank files are in Data S1.

#### Crosses to determine cleavage to LOF of TS-*ClvR^dbe^*

We crossed homozygous TS-*ClvR^dbe^* and *ClvR^dbe^* (control) males to *w^1118^* virgins to generate heterozygous offspring. Heterozygous TS-*ClvR^dbe^* (or *ClvR^dbe^* control) virgins were crossed to *w^1118^* males, incubated at a permissive temperature of 22° C, and the offspring was scored for the presence of the dominant TS-*ClvR^dbe^* marker. Results are shown in Table S3.

#### Analysis of escapers

From the experiment to determine cleavage to LOF described above, we recovered 91 males that did not carry the TS-*ClvR^dbe^* marker. We also recovered 72 males that did not carry the *ClvR^dbe^* marker from the control crosses with the original *ClvR^dbe^* flies. All of them were crossed to heterozygous TS-*ClvR^dbe^/+* (or *ClvR^dbe^* for the controls) females and incubated at 22° C again. After they mated, we took the male out of each vial and extracted genomic DNA. We amplified an amplicon spanning all 4 cut sites within the endogenous *dbe* locus and sequenced it. The offspring of the crosses was again scored for the presence of the TS *-ClvR^dbe^* (or *ClvR^dbe^*) marker. Afterwards, we selected 12 vials with low cleavage to LOF rates and transferred all the offspring to a food bottle to start a gene drive experiment as described below. In these gene drive experiments, we did not score marker frequencies. The drive experiment was continued until TS-*ClvR^dbe^* (or *ClvR^dbe^* controls) reached genotype fixation in all bottles. This took from 3 to 5 generations. Bottles with TS-*ClvR^dbe^* were subsequently transferred again and incubated at 29° C to test if a population collapse could be induced. All results with a more detailed description are shown in Data S1. The populations did crash, indicating that no functional endogenous alleles exist in these drive populations.

#### Crosses to test for temperature-dependent *Rescue* function of TS-*ClvR^dbe^*

We set up crosses involving females and males (all reared at 22° C) of the following genotypes: homozygous TS-*ClvR^dbe^* (10 vials), *w^1118^* (control, 5 vials), and *ClvR^dbe^* (control, 5 vials). These were incubated at a potentially restrictive temperature of 29° C. Offspring output of generations F1 and F2 are shown in Table S4.

#### Crosses to determine fecundity of TS-*ClvR^dbe^* flies over a range of temperatures

We set up 5 different crosses (genotypes below). These included 5 females and 5 males (4 replicates) that had been reared at 22° C. After setting up the cross, the vials were incubated at 23° C, 25° C, 27° C, and 29° C. Every 48 hours adults were transferred to a fresh food vial, and this was repeated 5 times. We scored the adult fly output in each of these vials. Results are shown in Fig. 2 and Fig. S2. Crosses were:

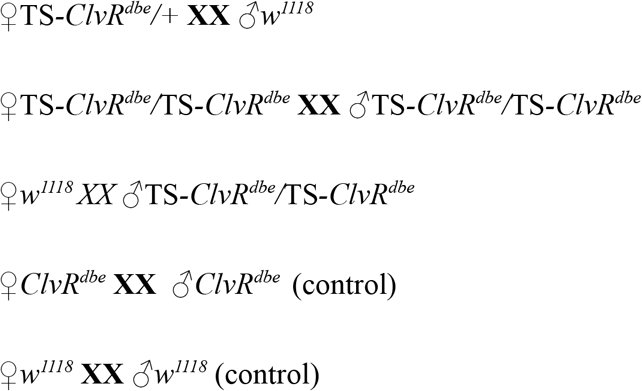

#### Gene drive experiment

We seeded 4 replicate populations by crossing heterozygous TS-*ClvR^dbe^*/+ males (or *Rescue*-INT^TS^/+ that do not have Cas9/gRNAs as a control) to *w^1118^* females (25% starting allele frequency). Flies were placed in food bottles, incubated at 22° C, and allowed to lay eggs for one day. Afterwards, they were removed from the bottles and the eggs were allowed to develop into adults. After approximately 16-17 days a large number had eclosed as adults. These were anesthetized on a CO_2_-pad, scored for the dominant TS-*ClvR^dbe^* marker, and transferred to a fresh food bottle to repeat the cycle. Counts are in Data S1.

#### TS-*ClvR^dbe^* and *ClvR^dbe^* (control) populations at 29°C

After the TS-*ClvR^dbe^* flies in the gene drive experiment reached genotype fixation (generation 10 and following), we first transferred them to a fresh food bottle to continue the gene drive experiment as described above. After they laid eggs in that bottle for one day, we transferred them again to a fresh bottle. That second bottle was now incubated at 29° C. Flies were given two days to lay eggs in that bottle before they were removed again. Eggs were allowed to develop into adults that were then scored and put in a fresh food bottle that was again kept at 29° C. Flies were kept in that bottle for one week prior to removal, so as to maximize the number of eggs laid. However, no progeny developed within these bottles. Results are shown in Fig. 3B (gray lines) and Table S6.

As a control experiment, we used the previously characterized *ClvR^dbe^* stock, which carries a WT copy of the recoded *Rescue* (19). *ClvR^dbe^* flies were taken from a gene drive experiment (generation 44, (19)), transferred to a fresh food bottle, and incubated alongside the TS-*ClvR^dbe^* bottles at 29° C. They were allowed to lay eggs for 2 days, after which adults were removed. After the eggs developed into adults, we determined the adult population number and transferred these individuals to a fresh food bottle to repeat the cycle. This was repeated for a total of 6 transfers with no obvious reduction in population size. Results are shown in Table S7.

**Fig. S1:**
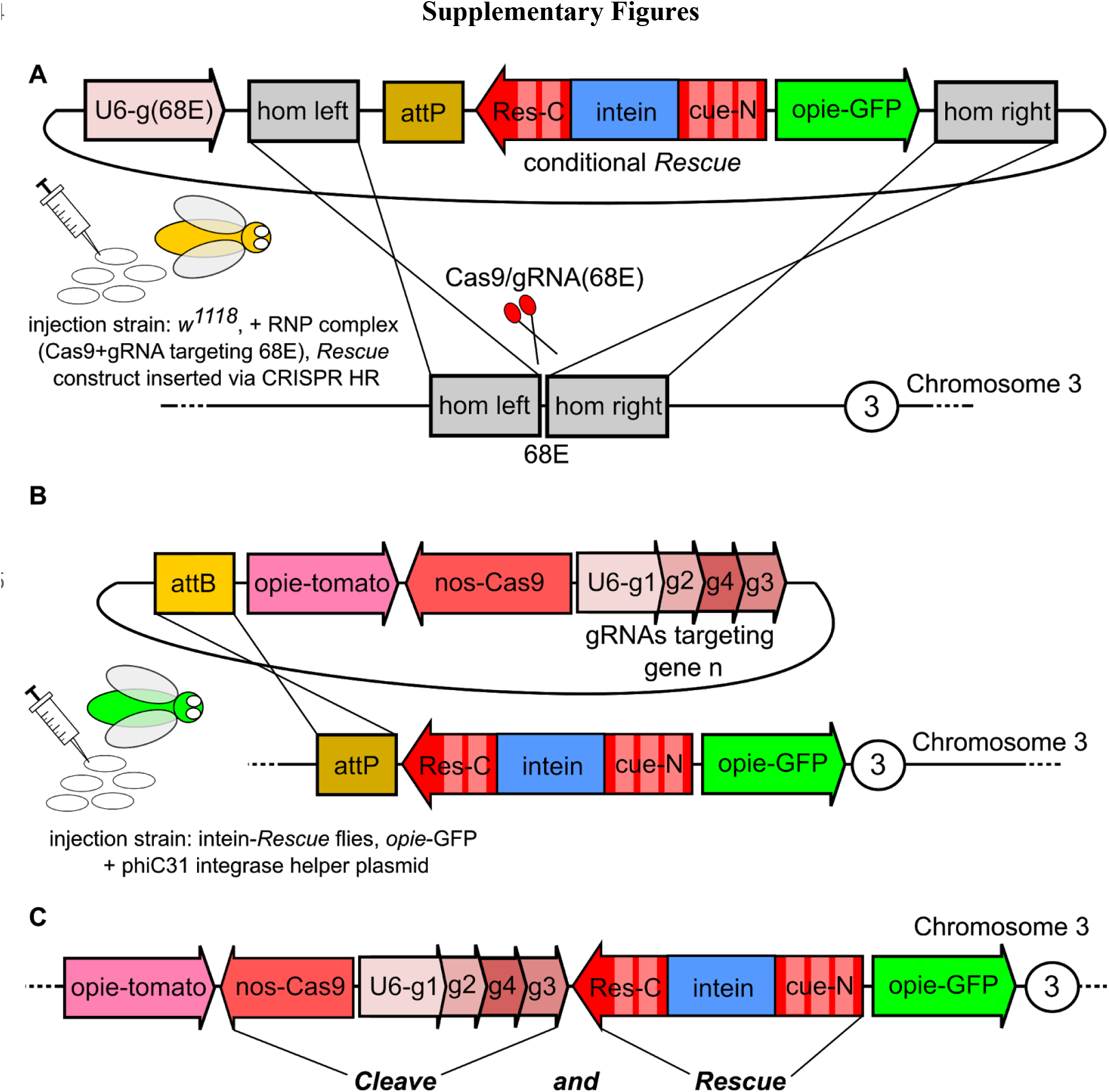
**(A) Genomic insertion of the *Rescue*-INT constructs.** We assembled plasmids that had TS and WT versions of the VMA intein inserted into the coding regions of *dbe* and *tko*, The constructs also had an ubiquitous *OpIE*-GFP marker, and an attP landing site for subsequent modifications of the locus. These were flanked by homology arms to facilitate CRISPR-HR mediated insertion into the genome. The construct was injected into *w^1118^* flies alongside a Cas9 RNP complex that targeted the genomic region at 68E on the third chromosome. **(B) Genomic integration of Cas9/gRNAs.** The second part of the *ClvR* drive mechanism, Cas9 and the gRNAs, were integrated into the genomic site of the TS-*Rescue* to yield complete TS-*ClvR^dbe^* flies. This second step was performed only with flies carrying the INT^TS^(*dbe*)Cys2. **(C) Schematic of the final TS-*ClvR^dbe^* drive element.**

**Fig. S2:**
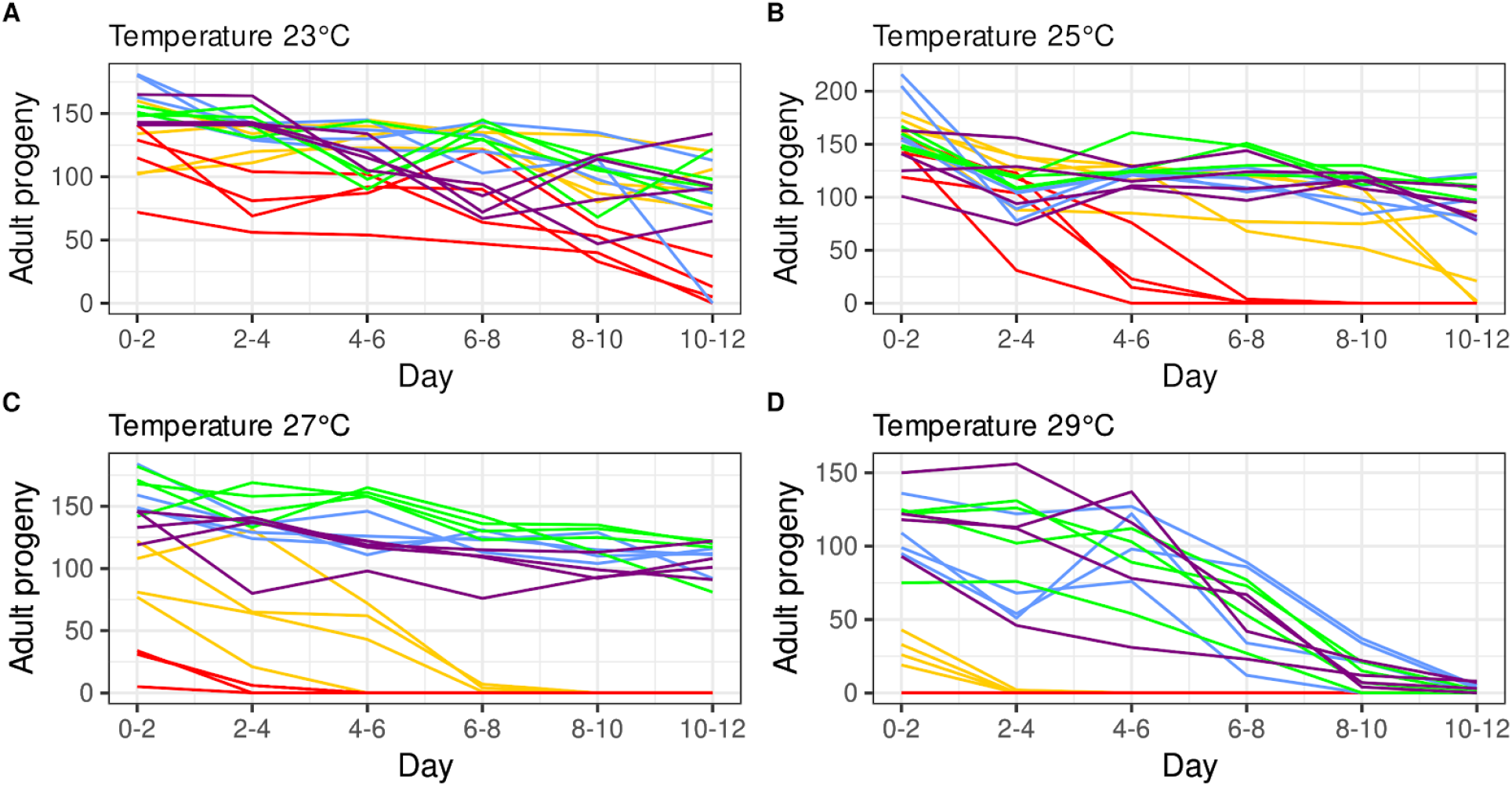
Adult fly output at different temperatures. Shown are the numbers of adult flies in four replicates that eclosed from different crosses incubated at 23° C **(A)**, 25° C **(B)**, 27° C **(C)**, and 29° C **(D)** over 12 days of egg-laying. Crosses were ♀TS-*ClvR^dbe^*/+ XX *♂w^1118^* in red, ♀TS-*ClvR^dbe^*/TS-*ClvR^dbe^* XX *♂*TS-*ClvR^dbe^*/TS-*ClvR^dbe^* in yellow, ♀*w^1118^ XX ♂*TS-*ClvR^dbe^*/TS-*ClvR^dbe^* in violet, ♀*ClvR^dbe^* XX *♂ClvR^dbe^* (control) in green, and ♀*w^1118^ XX ♂w^1118^* (control) in blue. Cumulative sums of adult progeny are shown in Fig. 2 in the main text.

**Fig. S3:**
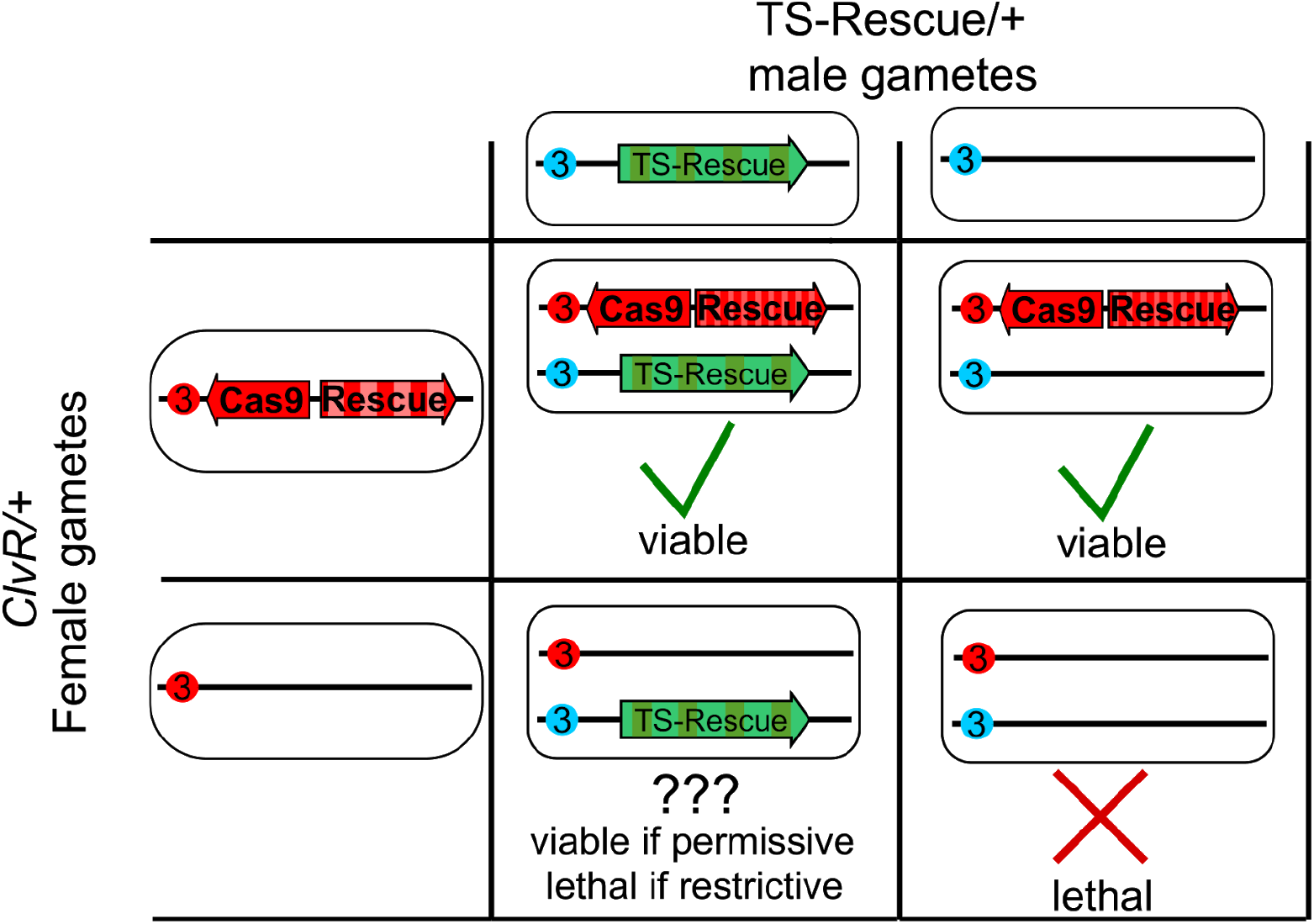
Crossing scheme to identify *conditional Rescue* candidates. The cross was set up with heterozygous *ClvR* females and heterozygous males carrying a single copy of the Rescue-INT^TS^. Cas9 and gRNAs that cleave the target gene render it LOF in the female germline and in the zygote due to maternal carryover-dependent cleavage of the paternal allele. The only functional copies of the target are provided by the *Rescue* in *ClvR* and/or the conditional *Rescue* in Rescue-INT^TS^. Half of the progeny, those that inherit the ClvR element, will always survive (upper row in Punnett square). Progeny that does not inherit *ClvR* or *Rescue*-INT^TS^ will always die if Cas9 cleaved the target (lower row right Punnett). In the cross we focused on flies that carried only the Rescue-INT^TS^ (lower row left Punnett) construct and are now in a background in which the endogenous version of the target gene has been rendered LOF. A good *Rescue*-INT^TS^ candidate should rescue viability at permissive low temperatures but not at restrictive high temperatures. Results of all screening crosses are in Table S1 and S2. Only flies carrying a TS-intein inserted after cysteine 2 of *dbe* showed the above behavior and were used to synthesize a full TS-*ClvR* element by integrating Cas9/gRNAs into that locus with PhiC31.

## Supplementary Tables

**Table S1:** Screening of *Rescue*-INT function for *tko.* Shown are the numbers of offspring from single fly crosses of heterozygous female *ClvR^tko^*/+ to males that carry a copy of different versions of the *Rescue-INT* for *tko*, Crosses were kept at 23° C or 27° C. WT (+/+) offspring were dying from maternal carryover activity of *ClvR^tko^* at both temperatures. Offspring that carry the *Rescue* within *ClvR^tko^* were not affected by temperature. Neither INT^WT^ nor INT^TS^ versions of the *tko Rescue* were able to rescue the LOF phenotypes induced by the *ClvR^tko^* element. Note: We did not obtain transformants for the INT^TS^ version inserted after cysteine 2 of *tko*. Since the INT^WT^ version of that *Rescue* did not provide gene function we reasoned that the TS-version will not work either. Thus, the construct was not pursued further.

| *Rescue* | Temperature | Replicate | *Rescue*/+ | *Rescue*/*ClvR* | *ClvR*/+ | +/+ | notes |
| --- | --- | --- | --- | --- | --- | --- | --- |
| INT<sup>TS</sup>(tko)Cys1 | 23 | A | 0 | 41 | 46 | 0 |  |
|  | 23 | B | 0 | 42 | 53 | 0 |  |
|  | 23 | C | 0 | 50 | 55 | 0 |  |
|  | 27 | A | 0 | 49 | 49 | 0 |  |
|  | 27 | B | 0 | 51 | 59 | 0 |  |
|  | 27 | C | 0 | 44 | 41 | 0 |  |
| INT<sup>TS</sup>(tko)Cys3 | 23 | A | 0 | 45 | 43 | 0 |  |
|  | 23 | B | 0 | 53 | 58 | 0 |  |
|  | 23 | C | 0 | 41 | 39 | 0 |  |
|  | 27 | A | 0 | 44 | 46 | 0 |  |
|  | 27 | B | 0 | 60 | 53 | 0 |  |
|  | 27 | C | 0 | 41 | 36 | 0 |  |
| INT<sup>WT</sup>(tko)Cys1 | 23 | A | 0 | 58 | 48 | 0 |  |
|  | 23 | B | 0 | 54 | 52 | 0 |  |
|  | 23 | C | 0 | 55 | 48 | 0 |  |
|  | 27 | A | 0 | 44 | 48 | 0 |  |
|  | 27 | B | 0 | 41 | 40 | 0 |  |
|  | 27 | C | - | - | - | - | sterile |
| INT<sup>WT</sup>(tko)Cys2 | 23 | A | 0 | 48 | 44 | 0 |  |
|  | 23 | B | 1 | 42 | 45 | 0 |  |
|  | 23 | C | 0 | 53 | 60 | 0 |  |
|  | 27 | A | 0 | 52 | 57 | 0 |  |
|  | 27 | B | 0 | 40 | 45 | 0 |  |
|  | 27 | C | 0 | 62 | 60 | 0 |  |
| INT<sup>WT</sup>(tko)Cys3 | 23 | A | 1 | 40 | 42 | 0 |  |
|  | 23 | B | 0 | 50 | 46 | 0 |  |
|  | 23 | C | 0 | 45 | 44 | 0 |  |
|  | 27 | A | 0 | 47 | 47 | 0 |  |
|  | 27 | B | 0 | 36 | 40 | 0 |  |
|  | 27 | C | 0 | 63 | 62 | 0 |  |

**Table S2.** Screening of intein-*Rescue* function for *dbe.* Shown are the numbers of adult offspring output from single fly crosses of heterozygous female *ClvR^dbe^* /+ to males that carry a copy of different versions of the *Rescue*-INT for *dbe*, Crosses were kept at 23° C or 27° C. WT (+/+) offspring of *ClvR^dbe^* mothers die due to LOF allele creation in the female germline and zygote at both temperatures. Offspring that carry the *Rescue* within *ClvR^dbe^* were not affected by temperature. Versions with a INT^WT^ inserted after cysteine 2 or 3 of *dbe* were functional at both temperatures. Versions with a INT^TS^ inserted after cysteine and 1 and 3 did not provide *Rescue* function at either temperature. However, a INT^TS^ inserted after cysteine 2 showed promising behavior, having *Rescue* activity at 23° C, whereas at 27° C most of the flies that carried it did not develop into adults (highlighted in red). We chose this *Rescue*-INT^TS^ to build a full TS-*ClvR* element by inserting Cas9 and gRNAs from *ClvR^dbe^*. Note: For the INT^WT^ inserted after cysteine 1 we did not obtain transformants after a first round of injections. Since the INT^TS^ version inserted after cysteine 1 did not show any *Rescue* activity we did not pursue this construct further.

| *Rescue* | Temperature | Replicate | *Rescue*/+ | *Rescue*/*ClvR* | *ClvR*/+ | +/+ |
| --- | --- | --- | --- | --- | --- | --- |
| INT<sup>TS</sup>(dbe)Cys1 | 23 | A | 0 | 35 | 39 | 0 |
|  | 23 | B | 0 | 35 | 32 | 0 |
|  | 23 | C | 0 | 31 | 36 | 0 |
|  | 27 | A | 0 | 40 | 45 | 0 |
|  | 27 | B | 0 | 39 | 32 | 0 |
|  | 27 | C | 0 | 47 | 50 | 0 |
| INT<sup>TS</sup>(dbe)Cys2 | 23 | A | 20 | 26 | 23 | 0 |
|  | 23 | B | 31 | 24 | 24 | 2 |
|  | 23 | C | 30 | 32 | 36 | 0 |
|  | 27 | A | 3 | 25 | 23 | 0 |
|  | 27 | B | 4 | 37 | 39 | 0 |
|  | 27 | C | 3 | 28 | 24 | 0 |
| INT<sup>TS</sup>(dbe)Cys3 | 23 | A | 0 | 49 | 49 | 0 |
|  | 23 | B | 0 | 33 | 35 | 0 |
|  | 23 | C | 0 | 36 | 32 | 0 |
|  | 27 | A | 0 | 44 | 47 | 0 |
|  | 27 | B | 0 | 38 | 37 | 0 |
|  | 27 | C | 0 | 32 | 30 | 0 |
| INT<sup>WT</sup>(dbe)Cys2 | 23 | A | 47 | 45 | 54 | 0 |
|  | 23 | B | 41 | 30 | 25 | 0 |
|  | 23 | C | 48 | 46 | 43 | 0 |
|  | 27 | A | 41 | 33 | 26 | 0 |
|  | 27 | B | 29 | 23 | 31 | 0 |
|  | 27 | C | 26 | 41 | 43 | 0 |
| INT<sup>WT</sup>(dbe)Cys3 | 23 | A | 47 | 43 | 52 | 0 |
|  | 23 | B | 20 | 52 | 40 | 0 |
|  | 23 | C | 6 | 51 | 36 | 0 |
|  | 27 | A | 41 | 33 | 26 | 0 |
|  | 27 | B | 29 | 23 | 31 | 0 |
|  | 27 | C | 26 | 41 | 43 | 0 |

**Table S3:** Cleavage to LOF of *ClvR^dbe^* and TS-*ClvR^dbe^* at 22°C. We assayed the cleavage activity of TS-*ClvR^dbe^* at the permissive temperature of 22° C by crossing heterozygous TS-*ClvR^dbe^* females to *w^1118^* males and scoring the offspring for the dominant *td-tomato* marker. The observed frequency of TS-*ClvR*-bearing flies in the offspring was lower than what we previously observed with *ClvR^dbe^* (>99%, (19)). That experiment was performed at a higher temperature of 26° C. Since the cleaving components (Cas9/gRNAs) of TS-*ClvR^dbe^* are exactly the same as for *ClvR^dbe^* we reasoned that the lower cleavage activity might be due to the lower incubation temperature. To confirm this, we set up the same crosses with the original *ClvR^dbe^* stock incubated at 22° C and found a lower rate of cleavage to LOF in that stock as well.

| Control crosses ♀ *ClvR<sup>dbe</sup>/+* XX ♂ *w<sup>1118</sup>* |  |  |  |  |  |  |
| --- | --- | --- | --- | --- | --- | --- |
| Bottle | *ClvR*-bearing | ♂ *w<sup>1118</sup>* | ♀ *w<sup>1118</sup>* | sum | *ClvR*-freq (%) | cleavage to LOF (%) |
| A | 817 | 17 | 14 | 848 | 96.34 | 92.69 |
| B | 800 | 21 | 19 | 840 | 95.24 | 90.48 |
| C | 831 | 16 | 13 | 860 | 96.63 | 93.26 |
| D | 597 | 18 | 14 | 629 | 94.91 | 89.83 |
| **total** | **3045** | **72** | **60** | **3177** | **95.85** | **91.69** |

| Crosses with TS-*ClvR<sup>dbe</sup>/+* XX *w<sup>1118</sup>* |  |  |  |  |  |  |
| --- | --- | --- | --- | --- | --- | --- |
| Bottle | *ClvR*-bearing | ♂ *w<sup>1118</sup>* | ♀ *w<sup>1118</sup>* | sum | *ClvR*-freq (%) | cleavage to LOF (%) |
| A | 832 | 22 | 24 | 878 | 94.76 | 89.52 |
| B | 975 | 31 | 37 | 1043 | 93.48 | 86.96 |
| C | 385 | 14 | 10 | 409 | 94.13 | 88.26 |
| D | 575 | 24 | 22 | 621 | 92.59 | 85.19 |
| **total** | **2767** | **91** | **93** | **2951** | **93.76** | **87.53** |

**Table S4.** Incubations at a restrictive temperature of 29°C. In a first test we crossed homozygous ♀TS-*ClvR^dbe^* to ♂TS-*ClvR^dbe^* and incubated them at a potentially restrictive temperature of 29° C. We also set up controls with *w^1118^* XX *w^1118^* and homozygous *ClvR^dbe^* XX *ClvR^dbe^*. All flies were reared at 22°C, crossed to each other in a fresh food vial and transferred to a 29° C incubator. All the crosses were fertile and gave progeny in the F1 generation. We transferred all the F1 flies to a fresh vial and kept them at 29° C. F1 progeny of TS-*ClvR^dbe^* XX TS-*ClvR^dbe^* was completely sterile, whereas F1 progeny from the two control crosses remained fertile. F1 progeny of all crosses was monitored for 1 week at 29° C. Afterwards we took two male TS-*ClvR^dbe^* flies and crossed them to *w^1118^* virgins. We also took two females and crossed them to *w^1118^* males. Both crosses did not yield offspring. The remaining F1 flies of the TS-*ClvR^dbe^* cross were put back at 22° C. And monitored for another week after which most of them had died. All of the flies remained sterile.

| Cross | Vial | F1 | F2 |
| --- | --- | --- | --- |
| *w<sup>1118</sup>* XX *w<sup>1118</sup>* | 1 | 66 | fertile |
|  | 2 | 85 | fertile |
|  | 3 | 102 | fertile |
|  | 4 | 110 | fertile |
|  | 5 | 95 | fertile |
| *ClvR<sup>dbe</sup>* XX *ClvR<sup>dbe</sup>* | 1 | 95 | fertile |
|  | 2 | 99 | fertile |
|  | 3 | 98 | fertile |
|  | 4 | 103 | fertile |
|  | 5 | 106 | fertile |
| TS-*ClvR<sup>dbe</sup>* XX TS-*ClvR<sup>dbe</sup>* | 1 | 32 | sterile |
|  | 2 | 60 | sterile |
|  | 3 | 63 | sterile |
|  | 4 | 25 | sterile |
|  | 5 | 18 | sterile |
|  | 6 | 64 | sterile |
|  | 7 | 69 | sterile |
|  | 8 | 67 | sterile |
|  | 9 | 63 | sterile |
|  | 10 | 83 | sterile |

**Table S5.** Allele frequencies of TS-*ClvR^dbe^* in the drive experiment at generation 18. Allele frequencies were measured by individually outcrossing 100 males from the drive populations to *w^1118^* females. Males that produced 100% TS-*ClvR* bearing offspring were considered to be homozygous. Males that produced 50% TS-*ClvR* bearing offspring were considered to be heterozygous.

| Drive replicate | homozygous | heterozygous | total alleles | allele freq (%)<br>TS-*ClvR* |
| --- | --- | --- | --- | --- |
| A | 71 | 4 | 150 | 97.33 |
| B | 70 | 11 | 162 | 93.21 |
| C | 61 | 6 | 134 | 95.52 |
| D | 78 | 4 | 164 | 97.56 |

**Table S6:** Incubation of gene drive populations at a restrictive temperature of 29° C. Flies from the gene drive experiment were transferred to a fresh food bottle and incubated at 29° C. They produced offspring for one more generation. That next generation was sterile resulting in a complete population collapse.

| Replicate | Generation drive | output in Generation n+1 | output in Generation n+2 |
| --- | --- | --- | --- |
| A | 10 | 486 | 0 |
| B | 10 | 575 | 0 |
| C | 10 | 433 | 0 |
| D | 10 | 188 | 0 |
| A | 12 | 513 | 0 |
| B | 12 | 477 | 0 |
| C | 12 | 430 | 0 |
| D | 12 | 367 | 0 |
| A | 13 | 476 | 0 |
| B | 13 | 481 | 0 |
| C | 13 | 588 | 0 |
| D | 13 | 456 | 0 |
| A | 16 | 321 | 0 |
| B | 16 | 272 | 0 |
| C | 16 | 251 | 0 |
| D | 16 | 369 | 0 |
| A | 17 | 245 | 0 |
| B | 17 | 295 | 0 |
| C | 17 | 214 | 0 |
| D | 17 | 229 | 0 |

**Table S7:** *ClvR^dbe^* drive populations at 29° C. As a control we took flies carrying *ClvR^dbe^* (non-TS) from a previously performed gene drive experiment (19) and transferred them to an incubator at 29° C. Flies were handled as with the other gene drive experiments. Every generation was transferred to a fresh food bottle and always kept at 29° C. This cycle was repeated for a total of 6 generations. Population size remained constant around the carrying capacity of the food bottles with no obvious fitness effects.

| Replicate | Generation drive (n) | Fly output generation (n+1) | Fly output generation (n+2) | Fly output generation (n+3) | Fly output generation (n+4) | Fly output generation (n+5) | Fly output generation (n+6) |
| --- | --- | --- | --- | --- | --- | --- | --- |
| A | 44 | 581 | 430 | 642 | 508 | 416 | 488 |
| B | 44 | 575 | 547 | 611 | 610 | 535 | 530 |
| C | 44 | 540 | 514 | 678 | 636 | 516 | 606 |
| D | 44 | 381 | 445 | 709 | 657 | 403 | 484 |

